# Crystallographic molecular replacement using an *in silico*-generated search model of SARS-CoV-2 ORF8

**DOI:** 10.1101/2021.01.05.425441

**Authors:** Thomas G. Flower, James H. Hurley

## Abstract

The majority of crystal structures are determined by the method of molecular replacement (MR). The range of application of MR is limited mainly by the need for an accurate search model. In most cases, pre-existing experimentally determined structures are used as search models. In favorable cases, *ab initio* predicted structures have yielded search models adequate for molecular replacement. The ORF8 protein of SARS-CoV-2 represents a challenging case for MR using an *ab initio* prediction because ORF8 has an all β-sheet fold and few orthologs. We previously determined experimentally the structure of ORF8 using the single anomalous dispersion (SAD) phasing method, having been unable to find an MR solution to the crystallographic phase problem. Following a report of an accurate prediction of the ORF8 structure, we assessed whether the predicted model would have succeeded as an MR search model. A phase problem solution was found, and the resulting structure was refined, yielding structural parameters equivalent to the original experimental solution.

## Introduction

Two key pieces of information are required to determine a macromolecular structure from a crystallographic diffraction experiment, namely the amplitudes and phases of the diffracted waves. The amplitudes of the diffracted waves are calculated directly from the measured intensities of the scattered waves, while the phase information is lost (1). The crystallographic phase problem continues remains a substantial bottleneck in macromolecular structure determination. In the majority of cases, the phase problem is overcome using an *in silico* method known as molecular replacement (MR) (2–5). In MR, a related structure, known as the search model, is used to provide initial phase-estimates for the target structure. While the advent of MR has rendered phase determination near-trivial in many cases, it was, until recently, limited to circumstances where structures of homologous proteins already existed.

In principle, advances in *in silico* protein structure prediction could provide molecular replacement search models sufficiently similar to the target structures so as to produce a MR phase solution (6–8). Such an advance could, in theory, forever bypass the need for experimentally derived phases. The use of *ab initio* models as MR search models has seen some success, principally with α helix-rich proteins (9). Accurate *ab initio* prediction of β-rich folds has long proven to be exceptionally challenging (10) due, in part, to the high proportion of nonlocal interactions between β-strands. To our knowledge only one β-rich crystal structure has previously been phased using an *ab initio* generated search model (11). This approach relied on a highly parallelized trial and error approach where hundreds of input MR ensemble search models were tested (10).

Recent advances in *ab initio* modelling have come from the long-accepted principle that evolutionary covariance of residues can aid in inter-residue contact prediction. This relies on the fact that deleterious point mutations are often paired with compensatory ones during evolutionary development. Multiple sequence alignments (MSA) of many related protein sequences can be used to identify these correlated mutations, with several systems applying neural networks to do so (12–16). Many groups, including Google DeepMind’s ‘AlphaFold’, have taken this principal one step further by predicting specified distances between residue pairs which provide more information about the structure than contact predictions alone (16–20). In the case of AlphaFold, these inter-residue distances, along with backbone torsion angles, are predicted in a first step using convolutional neural networks (16). They are then provided as a target in gradient descent algorithm, which aims to bring the 3D structure as close as possible to these predicted distance and torsion angles. These principles were applied to great effect during the 14^th^ biennial Critical Assessment of protein Structure Prediction (CASP) competition, where the AlphaFold2 team contributed a wealth of *ab initio* predictions, the majority of which were considered highly similar to experimentally determined protein structures with a median Global Distance Test (GDT) score of 92.4 out of 100 (21, 22).

Our group recently determined the crystal structure of ORF8 at 2.04 Å (23), a SARS-CoV-2 protein which has been implicated in immune evasion. The structure revealed that the protein is composed entirely of β-strands and unstructured regions. While a full native dataset was collected during the early stages of the project, a lack of suitable search model made phase determination by MR unfeasible. Initial phases were obtained by anomalous dispersion.

Here we assessed whether a template-free, *ab initio* protein model, generated by the AlphaFold2 group, was of sufficient quality to phase the native ORF8 dataset by MR. No truncation of the model was required nor was there any need to provide an ensemble of search models. Not only is this approach likely to prove useful for future structural determination campaigns where a homologous structure is not available but could aid in the determination of preexisting ‘unsolvable’ datasets.

## Results

Superposition of the *ab initio* AlphaFold2 ORF8 model (CASP14 model ID: T1064TS427_1-D1) with the previously deposited crystal structure protomer (PDBID: 7JTL) showed they are highly similar, exhibiting RMSDs of 0.68 Å and 0.92 Å with chains A and B respectively (Fig. 1). This suggests the potential of the AlphaFold2 model as an appropriate MR search model candidate.

**Figure 1.**
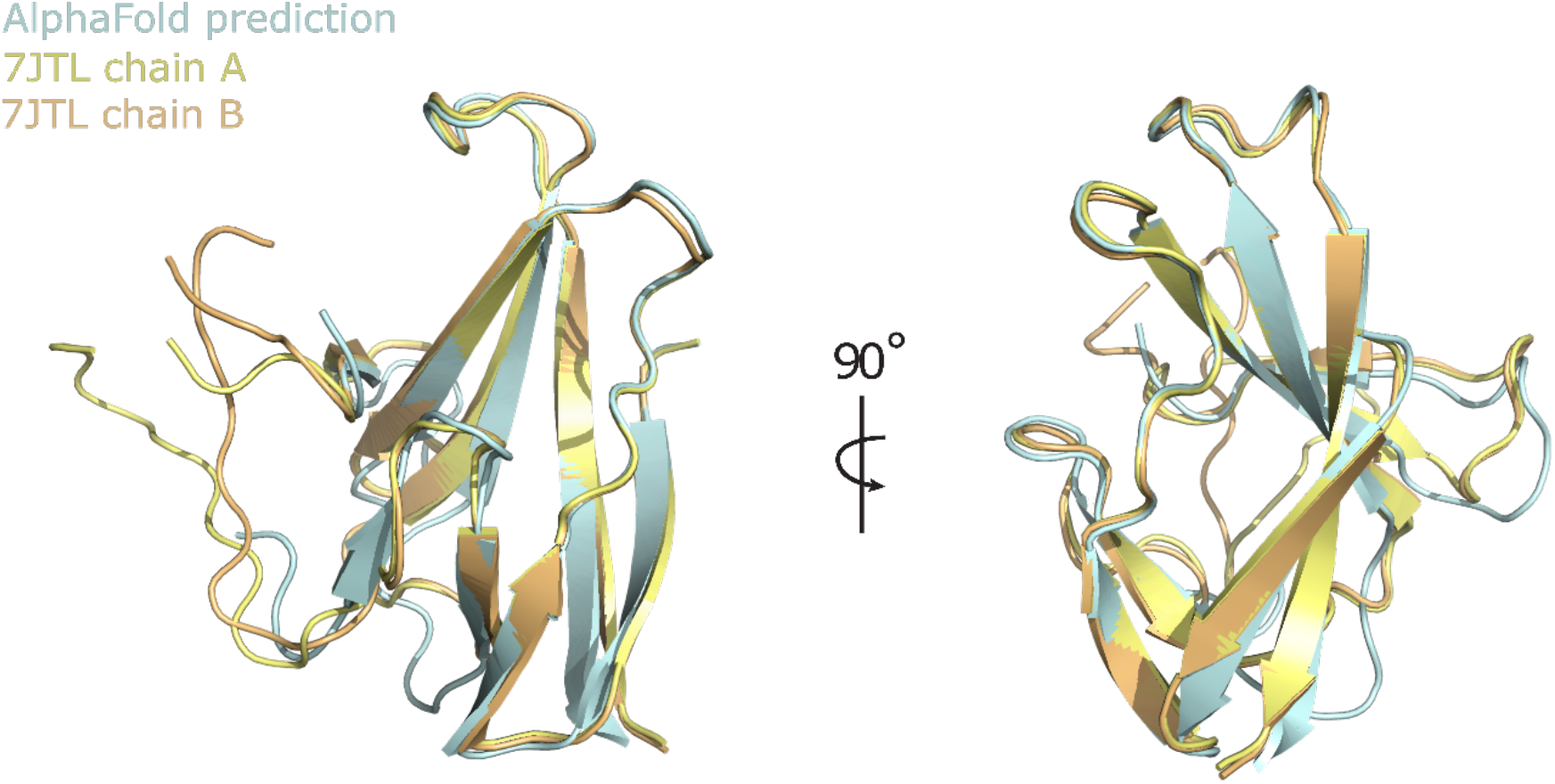
Superposition of experimentally determined CoV-2 ORF8 structure with AlphaFold2 prediction. Chains A and B of the ORF8 crystal structure (PDBID: 7JTL) are superposed and colored yellow and orange respectively. The AlphaFold2 prediction is colored cyan.

To test this, MR was performed using native ORF8 diffraction data/structure factor amplitudes (PDBID: 7JTL) and the unaltered AlphaFold2 ORF8 prediction as a search model. A single MR solution was identified with 2 copies in the asymmetric unit. The Log Likelihood Gain (LLG) was 167 with values of 40 or greater typically indicating a correct solution (24). The MR solution places the search model within the crystal lattice in near exact superposition with the asymmetric unit of the experimentally determined structure, confirming the search models were correctly placed (Fig. 2).

**Figure 2.**
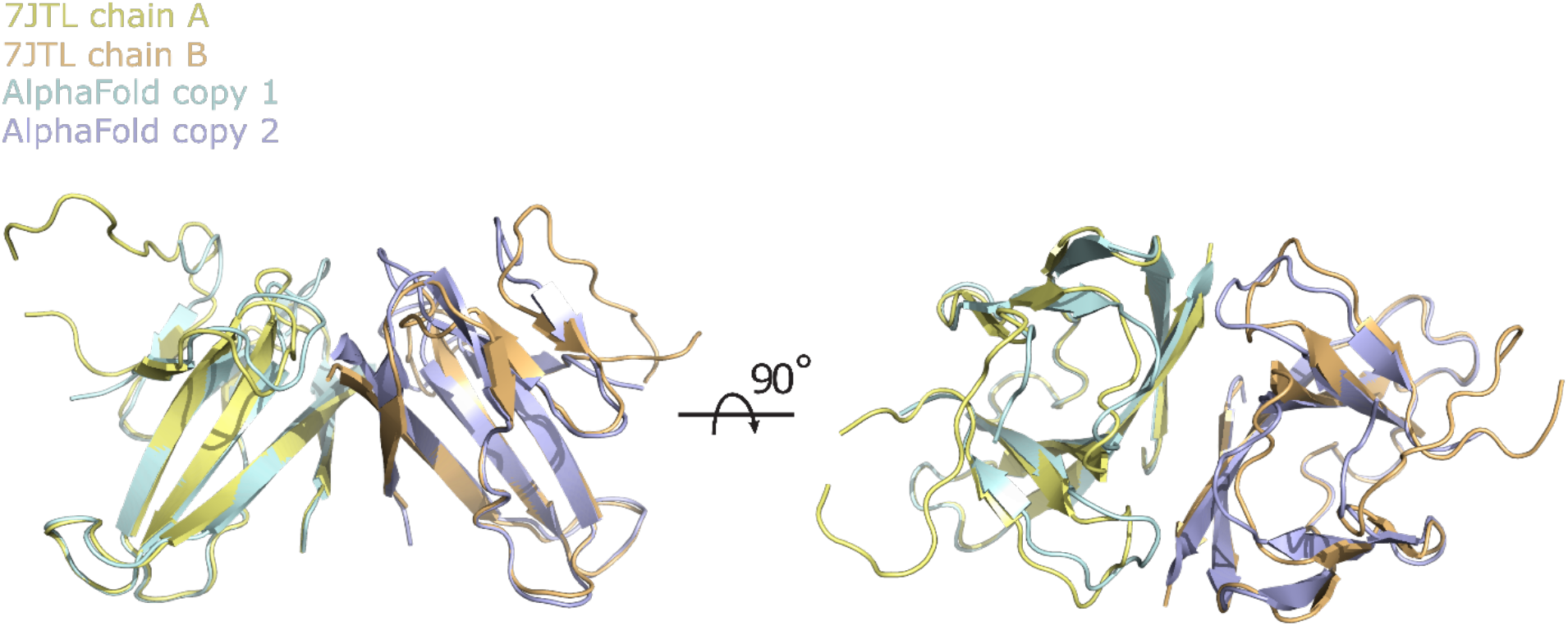
Position of two placed CoV-2 ORF8 AlphaFold2 search models following MR. MR was performed using native ORF8 diffraction data. The first and second placed copies of the search model are colored cyan and purple respectively. The positions of chains A and B of the experimentally-phased/previously determined ORF8 crystal structure (PDBID: 7JTL) asymmetric unit are shown for reference and are colored yellow and orange respectively.

A 2Fo-Fc electron density map was generated using phases from the placed but un-modified AlphaFold model and the native structure factor amplitudes (Fig. 3). The density was of sufficient quality to allow the majority of sidechains and unmodelled main chain regions to be built unambiguously. A number of map features were present that were in poor agreement with the input model (Fig. 3b). These differences exclude the possibility that the density is dominated by input model-based phase-bias. Such differences include incorrect side-chain positioning/orientation, minor main chain deviations and the presence of unbuilt/unmodelled regions. Iterative rounds of manual model building and refinement produced a final structure determination with global quality metrics on par with those of the previously deposited structure (Table 1).

**Figure 3.**
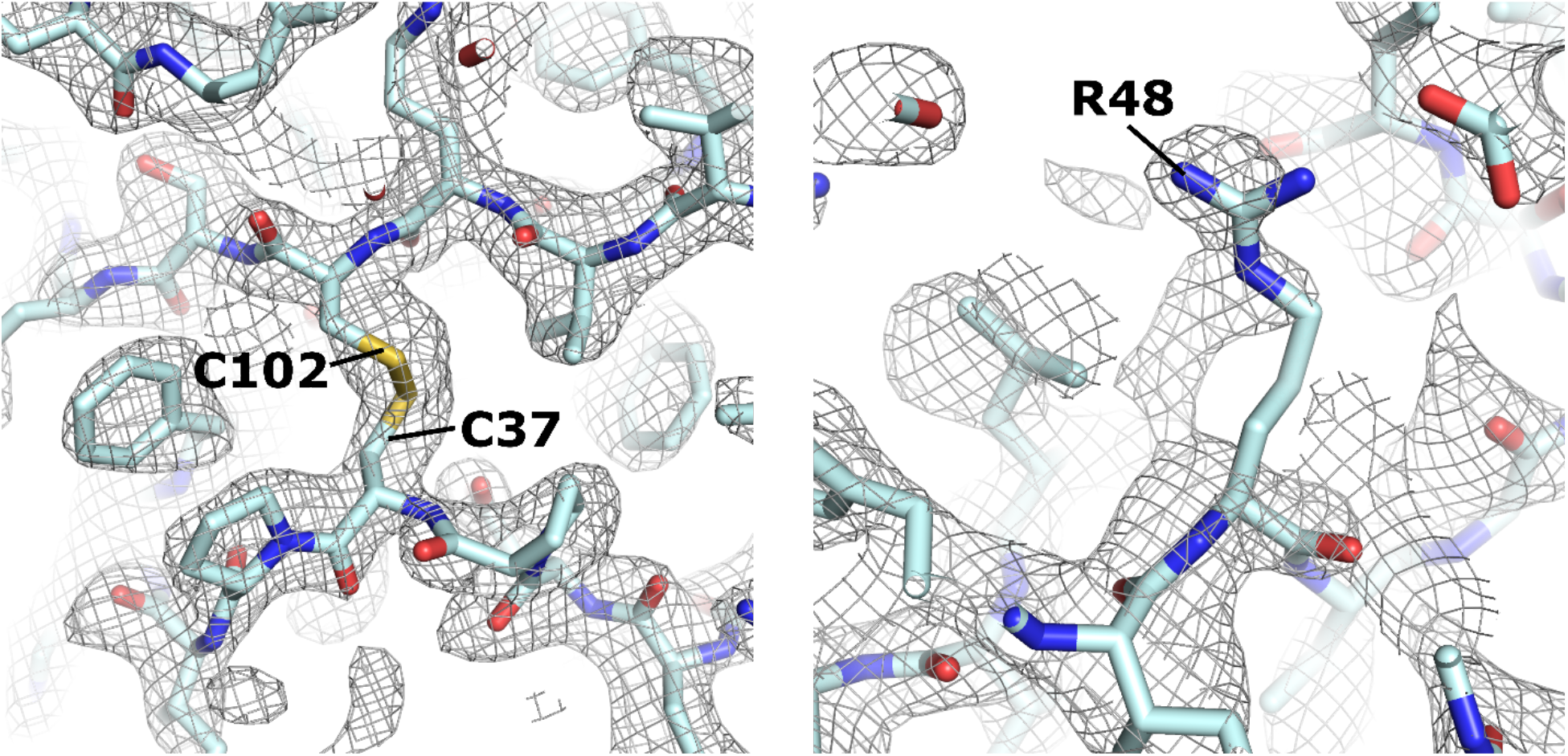
Representative regions of 2Fo-Fc electron density following MR with AlphaFold2 CoV-2 ORF8 search model. Left panel shows a region of density that is in good agreement with the unmodified AlphaFold2 search model. Right panel provides an example where there is an obvious discrepancy between the experimental data and the search model, suggesting that the position and orientation of the Arg48 side-chain should be modified. Map is contoured at 1.2 σ and represented as a grey mesh.

**Table 1.** Data collection and refinement statistics

|  | SARS-CoV-2 ORF8 |  |
| --- | --- | --- |
|  | Experimentally phased | AlphaFold2 MR |
| **Data collection** |  |  |
| Space group | $P4_12_12$ | $P4_12_12$ |
| Cell dimensions |  |  |
| $a, b, c$ (Å) | 44.3 44.3 264.1 | 44.3 44.3 264.1 |
| $\alpha, \beta, \gamma$ (°) | 90, 90, 90 | 90, 90, 90 |
| Resolution (Å) | 43.65-2.04 (2.113-2.04) | 43.65-2.04 (2.113-2.04) |
| $R_{\text{pim}}$ | 0.032 (0.555) | 0.032 (0.555) |
| $I / \sigma I$ | 14.92 (1.20) | 14.92 (1.20) |
| Completeness (%) | 97.8 (90.2) | 97.8 (90.2) |
| Redundancy | 10.0 (7.6) | 10.0 (7.6) |
| **Refinement** |  |  |
| Resolution (Å) | 43.65-2.04 | 43.65-2.04 |
| No. reflections | 17005 (1399) | 17005 (1399) |
| $R_{\text{work}} / R_{\text{free}}$ (%) | 22.0 (35.2)/ 26.4 (38.4) | 22.3 (35.4)/27.5 (42.2) |
| No. atoms |  |  |
| Protein | 1609 | 1625 |
| Water | 201 | 96 |
| $B$-factors | | |
| Protein | 45.4 | 44.3 |
| Water | 44.3 | 45.7 |
| R.m.s. deviations |  |  |
| Bond lengths (Å) | 0.008 | 0.010 |
| Bond angles (°) | 1.00 | 1.30 |

Values in parentheses are for highest-resolution shell

## Discussion

The success of the AlphaFold2 predictions at CASP14 has led to considerable excitement and discussion of the relative roles of *ab initio* prediction and experimental structure determination in the future (22). Our assessment found that the AlphaFold2 predicted structure of SARS-CoV-2 ORF8 was of sufficient quality to yield a correct MR solution. This represents both a stringent test of accuracy and bodes well for the practical utility of AlphaFold2 predictions in MR. It will clearly be important for the AlphaFold2 method, and the computing power needed to support it, to become available to the structural biology community.

Despite the impressive accuracy of the fold assignment in the AlphaFold prediction, critical details were missing and could only be resolved by crystallographic structure solution. In general, side-chain conformations were inaccurate in the predicted structure. Notably, dimerization of ORF8 is thought to be important for function, yet the AlphaFold2 prediction only provided the structure of the monomer. Even before AlphaFold2, high quality *ab initio* structure predictions were accurate enough to reduce, if not eliminate, the need for experimental determinations of the overall fold of single domain proteins (25). In our view, the real utility of the “accuracy revolution” in structural biology will be to increase synergy with experimentation. Not only will this be invaluable for crystallographic MR, as assessed here, but in the interpretation of density in cryo-EM as well.

## Acknowledgements

We thank Marc Allaire, Cosmo Buffalo, and Richard Hooy for helpful discussions and comments on the manuscript. This work was supported by UCOP emergency seed grant R00RG2347 (JHH), and National Institutes of Health grant R37 AI112442 (JHH).

## Methods

### Molecular Replacement

The AlphaFold2 ORF8 prediction was obtained from the CASP14 website (www.predictioncenter.org; model ID: T1064TS427_1-D1) and prepared for Molecular Replacement using the *Phenix* software suite (26). Molecular replacement was performed using the program *Phaser* with default search parameters. The prepared AlphaFold2 ORF8 prediction was provided as a search model and native ORF8 structure factors as experimental data. The search was limited to the known space group *P*4_1_2_1_2. A single solution was obtained with 2 copies in the asymmetric unit, an LLG value of 167 and TFZ score of 14.7 indicating that the correct solution had been found.

